# Quantification of microRNA editing using two-tailed RT-qPCR for improved biomarker discovery

**DOI:** 10.1101/2021.06.15.448409

**Authors:** Gjendine Voss, Anders Edsjö, Anders Bjartell, Yvonne Ceder

## Abstract

Even though microRNAs have been viewed as promising biomarkers for years, their clinical implementation is still lagging far behind. This is in part due to the lack of RT-qPCR technologies that can differentiate between microRNA isoforms. For example, A-to-I editing of microRNAs through adenosine deaminase acting on RNA (ADAR) enzymes can affect their expression levels and functional roles, but editing isoform-specific assays are not commercially available. Here, we describe RT-qPCR assays that are specific for editing isoforms, using microRNA-379 (miR-379) as a model. The assays are based on two-tailed RT-qPCR, and we show them to be compatible both with SYBR Green and hydrolysis-based chemistries, as well as with both qPCR and digital PCR. The assays could readily detect different miR-379 editing isoforms in various human tissues. We found that the miR-379 editing frequency was higher in prostate cancer samples compared to benign prostatic hyperplasia samples. Furthermore, decreased expression of unedited miR-379, but not edited miR-379, was associated with treatment resistance, metastasis and shorter overall survival. Taken together, this study presents the first RT-qPCR assays that were demonstrated to distinguish A-to-I-edited microRNAs, and shows that they can be useful in the identification of biomarkers that previously have been masked by other isoforms.

**Graphical abstract:** 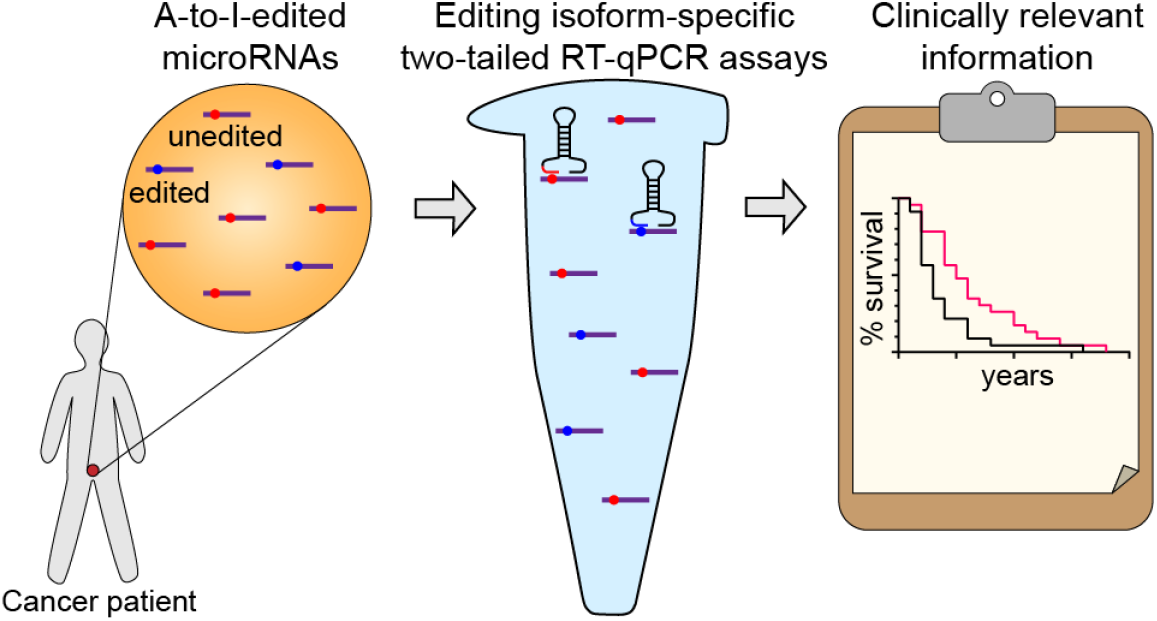

## Introduction

Ever since their discovery in 1993 (Lee et al. 1993, Wightman et al. 1993), microRNAs (miRNAs) have been a topic of interest in both basic research and in various disease contexts. These small non-coding RNAs act through binding different mRNA targets based on imperfect sequence complementarity, and regulate both mRNA stability and translation (Pasquinelli 2012). Typically, one miRNA has hundreds of different targets in a cell, which it binds with varying affinity. This allows miRNAs to regulate several players in the same pathway and multiple biological processes at once. As such, miRNAs are thought to be crucial for the maintenance of homeostasis in a cell. It is therefore not surprising that the deregulation of miRNAs is associated with several disease states (Fabris et al. 2016, Wendt et al. 2018, Fasolo et al. 2019, van den Berg et al. 2020), which has sparked interest in their suitability both as therapeutic targets or agents and as biomarkers. The use of miRNAs as biomarkers is supported by their remarkable stability in biological fluids (Mitchell et al. 2008) and even in harshly treated specimens such as formalin-fixed tissues for several years (Hui et al. 2009). However, even though there are thousands of studies describing the association between the expression of certain miRNAs and clinical parameters, there are currently no FDA-approved miRNA biomarkers, and only few that are ready to be used in clinical practice (Bonneau et al. 2019).

One possible reason for this discrepancy is the fact that miRNAs do not exist in only one static form, but actually occur in multiple different isoforms, which can differ in stability and carry out distinct functions in the cell. These different isoforms are produced by post-transcriptional modifications such as addition or removal of terminal nucleotides (isomiRs) and A-to-I editing of internal bases (Ameres and Zamore 2013).

A-to-I editing is the deamination of adenosine nucleotides to form inosine, carried out by adenosine deaminase acting on RNA (ADAR) enzymes. ADARs can target virtually any double-stranded RNA, including primary miRNAs (pri-miRNAs). As inosine preferentially base pairs with cytosine, A-to-I editing can alter the secondary structure of pri-miRNAs, leading to an inhibition of processing and maturation, which ultimately affects expression levels. If the seed sequence is altered, this can also cause the miRNA to bind a different set of mRNA targets (Kawahara et al. 2007, Shoshan et al. 2015, Nishikura 2016, Velazquez-Torres et al. 2018, Xu et al. 2019, van der Kwast et al. 2020).

Currently, the only technology to distinguish these miRNA editing isoforms is RNA sequencing, which is time-consuming, expensive, and requires high input to detect miRNAs with low expression. Additionally, library preparation can introduce bias, making it difficult to quantify relative abundance of miRNAs accurately (Witwer and Halushka 2016).

In contrast, commonly used quantitative PCR (qPCR) methods are not specific enough to distinguish individual nucleotide differences (Androvic et al. 2017). Therefore, if only one isoform is biologically relevant in certain contexts, but other isoforms of the same miRNA are detected as well, this could mask the deregulation of the clinically relevant miRNA.

For example, recent publications have demonstrated that the ratio of isomiRs differs between cell activation states (Nejad et al. 2018b, Pillman et al. 2019), and that conventional qPCR methods are often biased and may skew the results (Wu et al. 2007, Schamberger and Orbán 2014, Magee et al. 2017, Nejad et al. 2018b). They have also shown that polyadenylation-dependent RT-qPCR protocols can potentially be adaptable for accurate isomiR quantification (Nejad et al. 2018a). Another method that has made progress towards the accurate quantification of terminal isoforms is Dumbbell-PCR (Honda and Kirino 2015). Yet, these methods do not address internal A-to-I editing. A-to-I editing-sensitive qPCR protocols have been published for mRNAs (Chen et al. 2008), but they cannot be directly transferred for miRNAs, which are much shorter. For A-to-I-edited miRNAs, no commercial assays are available, and no validated methods have been published. Studies have attempted to quantify edited and unedited mature miRNAs with custom-ordered TaqMan assays (van der Kwast et al. 2018, van der Kwast et al. 2020). However, they only tested the amplification efficiency of the assays using serial dilutions of a biological sample, but provided no data regarding the cross-detection between editing isoforms. No validation using synthetic oligonucleotides to investigate amplification of each isoform on its own was carried out.

Hence, there is a dire need for an extensively validated RT-qPCR setup that can be proven to reliably distinguish A-to-I editing miRNA isoforms. If qPCR methods can be adapted and evaluated accordingly, this could potentially improve existing miRNA biomarkers, or reveal undiscovered candidates that depend on the editing status of the miRNA.

A disease that could benefit from refined biomarkers is prostate cancer (PC), which is often curable if detected early. However, more aggressive forms do exist with a rapid progression to metastatic disease, leaving no curative treatment options and ultimately resulting in the patient’s death. Finding biomarkers that early on can predict which patients are likely to develop aggressive disease and will require harsh treatment is therefore highly desirable in PC.

We have previously found that PC bone metastasis may be promoted by downregulation of microRNA-379 (miR-379; data submitted for publication), a miRNA that is known to be edited by ADAR2 at nucleotide 5 of mature miR-379-5p (Kawahara et al. 2008). The position of the edited nucleotide close to the Drosha processing site and in the seed sequence leads to both a decrease in processing efficiency (Kawahara et al. 2008) and binding of a different set of mRNA targets (Xu et al. 2019). The latter suggests that miR-379 editing isoforms may have distinct biological functions, but for lack of suitable assays, it has not been possible to pinpoint whether the anti-metastatic effect of miR-379 in PC is isoform-specific. We therefore chose miR-379 as a model to develop a method for the quantification of A-to-I-edited miRNA isoforms based on the recently described highly specific two-tailed RT-qPCR assays (Androvic et al. 2017). The principle of two-tailed RT-qPCR relies on using two short hemiprobes that bind cooperatively to prime reverse transcription rather than one longer RT primer. This greatly increases the specificity, as a single nucleotide mismatch will affect a short hemiprobe more than a long primer.

The assays developed by us were highly specific for individual editing isoforms of miR-379. Using a PC cohort, we found that the miR-379 editing frequency was higher in cancer samples compared to benign samples, and that low expression of the unedited miR-379 isoform was associated with metastasis, treatment resistance and shorter overall survival. This demonstrates that isoform-specific analysis of miRNA expression may reveal more clinical information than has been possible with previously available qPCR assays.

## Results and Discussion

### Sensitive and isoform-specific detection of A-to-I-edited miRNAs with two-tailed RT-qPCR

To test commercially available qPCR reagents for miR-379, we used TaqMan Advanced microRNA assays for miR-379 on dilutions of unedited and edited miR-379 molecules. We expected that they would either be specific for unedited miR-379 or recognize both isoforms indiscriminately of editing status. Instead, the assays detected both isoforms, but at vastly different PCR efficiencies (109% for unedited miR-379, 73% for edited miR-379), creating a different preference for one or the other isoform depending on the concentration (Figure 1). This makes this assay neither suitable for distinguishing editing isoforms, nor to quantify total miR-379 reliably. In addition, the tested assay required an input of 10^5^ molecules of miR-379 to detect the miRNA.

**Figure 1.**
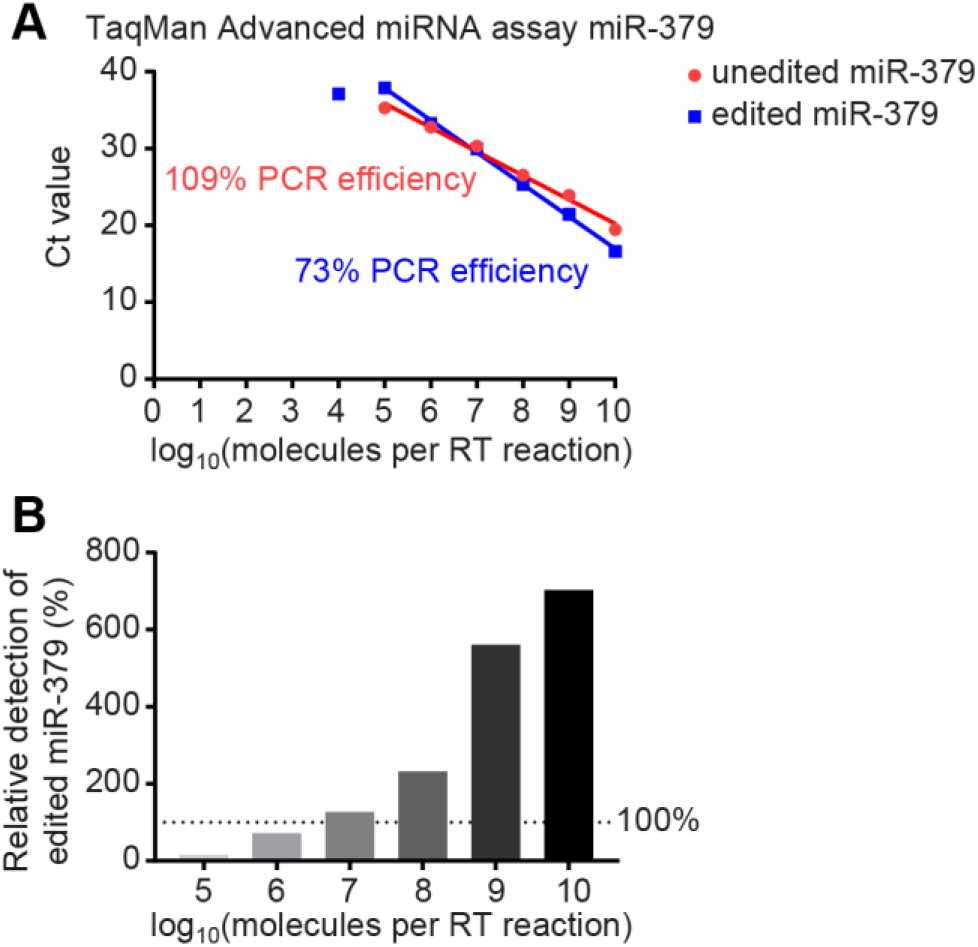
Detection of miR-379 editing isoforms with commercial TaqMan assays. (**A**) Ct values for different dilutions of unedited (red circles) and edited (blue squares) miR-379 RNA oligonucleotides quantified with TaqMan Advanced microRNA assays. PCR efficiencies were calculated based on the slope. Standard deviations were too small to be plotted as error bars. (**B**) Relative detection of edited miR-379 compared to an equal number of unedited miR-379 molecules as calculated from the difference in Ct cycles in (A).

In order to achieve editing isoform-specific reverse transcription (RT) and qPCR, we designed two-tailed RT primers as previously described (Androvic et al. 2017), placing the 5′ hemiprobe so that it would cover the editing site (Figure 2A). We tested different primer design parameters to get optimal specificity and sensitivity, trying different combinations of hemiprobe length and hemiprobe placement in relation to the edited nucleotide (Supplementary Table 4). A length of 5 nt was optimal for the 3′ hemiprobe (4 nt could not efficiently prime reverse transcription; 6 nt primed reverse transcription even of the non-target isoform, resulting in lower specificity). We found that the position of the edited nucleotide in the 5′ hemiprobe did not affect the specificity much, and ultimately we selected the probe with the highest sensitivity.

**Figure 2.**
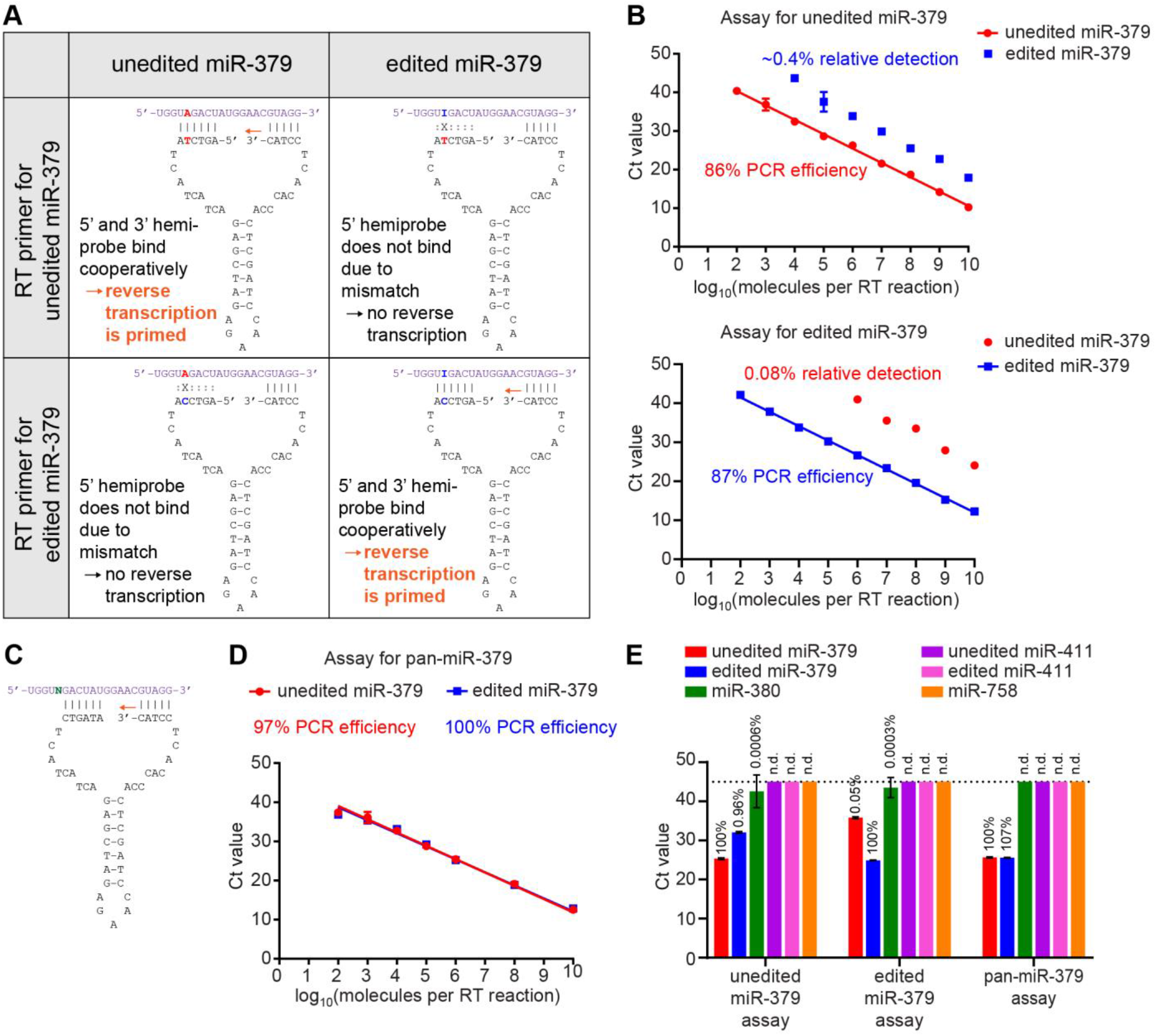
Two-tailed RT-qPCR assays for unedited, edited or pan-miR-379. (**A**) Principle of RT primer design for editing-sensitive miR-379 assays. The 5′ hemiprobe is designed to only bind one of the two editing isoforms. Reverse transcription is only primed when both hemiprobes can bind with high affinity. (**B**) Ct values for different dilutions of unedited (red circles) and edited (blue squares) miR-379 RNA oligonucleotides quantified by the two-tailed RT-qPCR assays specific for unedited miR-379 (top) and edited miR-379 (bottom). Error bars denote standard deviation; for points without error bars, the standard deviation was too small to be plotted. PCR efficiencies were calculated based on the slope. The average relative detection rate of non-target miR-379 was calculated based on the Ct difference to target miR-379. (**C**) RT primer design for the pan-miR-379 assay. The 5′ hemiprobe binds a part of the miRNA that is not subject to editing. (**D**) Ct values for different dilutions of unedited (red circles) and edited (blue squares) miR-379 RNA oligonucleotides quantified by the pan-miR- 379 RT-qPCR assay. Error bars denote standard deviation; for points without error bars, the standard deviation was too small to be plotted. PCR efficiencies were calculated based on the slope. (**E**) Ct values for 10^6^ molecules of different miR-379 family members by the three two-tailed RT-qPCR assays. Error bars denote standard deviation. The table lists the relative detection of each miRNA compared to the detection of the assay’s target (100%). The dotted line marks the maximum number of Ct cycles (45 cycles). Any samples that did not show amplification at a Ct lower than this are considered undetectable. n.d. = not detected

RT of dilutions of miR-379 RNA oligonucleotides with the two-tailed primers and subsequent SYBR Green qPCR showed that the designed probes could reliably detect their target isoforms over a large dynamic range with linearity between 100 and 10^10^ molecules per reaction (Figure 2B) and PCR efficiencies of 85–90%. Our RT-qPCR setting was 1000-fold more sensitive than the commercially available assay. Furthermore, the assays were very specific for their respective target isoform of miR-379 with relative detection of edited miR-379 by the unedited primers below 1%, and relative detection of unedited miR-379 by the edited primers below 0.1% (Figure 2B). To mimic a more complex mixture of RNA molecules, the same experiments were performed with the miR-379 molecules being diluted in yeast RNA, showing largely the same result (Supplementary Figure 1).

In certain contexts, it may be desirable to quantify all isoforms of a miRNA, independently of the editing status. For this purpose, we designed a pan-miR-379 RT primer with the 5′ hemiprobe shifted to bind outside of the editing site (Figure 2C). As the reverse primers in the qPCR are designed to bind the RT-extended part of the RT primer, *i.e.*, the part that is complementary to the miRNA (Androvic et al. 2017), the reverse primers inevitably have slightly different affinities to different isoforms. We found a 60/40 mixture of edited-specific and unedited-specific reverse qPCR primers to be optimal to amplify both miR-379 isoforms equally well without bias towards one or the other isoform (Figure 2D, Supplementary Figure 2). The ratio of detection of the two isoforms was close to 1.0 across all tested miR-379 concentrations, and the PCR efficiencies were 95–100% for both isoforms, implying that amplification of miR-379 with the pan-miR-379 primers is truly independent of editing status. Finally, we wanted to ensure that in order to be able to quantify miR-379 isoforms correctly in biological samples, other closely related members of the miR-379 family (Seitz et al. 2004) would not be amplified and skew the results. The miR-379-specific RT-qPCR primers did not efficiently reverse transcribe or amplify any of the miR-379 family members, with relative detection of miR-380 below 0.001%, and no detection of miR-411 or miR-758 (Figure 2E).

### Adaptation for hydrolysis probe-based qPCR and digital PCR

While SYBR Green-based two-tailed RT-qPCR is cost-efficient, there are applications for which one may want to utilize hydrolysis probes and/or digital PCR (dPCR). The distance between the forward and reverse qPCR primers allows for a 24 nt-long hydrolysis probe to be fit with a sufficiently high melting temperature and a 1–2 nt gap in between the primer and the hydrolysis probe to enable efficient extension and qPCR. As the hydrolysis probe targets part of the 3′ hemiprobe and the stem-loop sequence of the original RT primer, *i.e.* a sequence that is the same for both the unedited and the edited miR-379 assay, the same hydrolysis probe was used for both isoforms. Two-tailed RT-qPCR using hydrolysis probes worked well for both RT primers, with sensitivity down to 1000 molecules (Figure 3A). However, the relative detection of the non-target miR-379 was higher than with SYBR Green, with 2% detection of edited miR-379 in the unedited miR-379-specific setting, and 0.5% detection of unedited miR-379 in the edited miR-379-specific setting. The slight decrease in both sensitivity and specificity of hydrolysis-based qPCR can potentially be outweighed by the possibility to multiplex using several fluorophores in parallel for specific applications.

**Figure 3.**
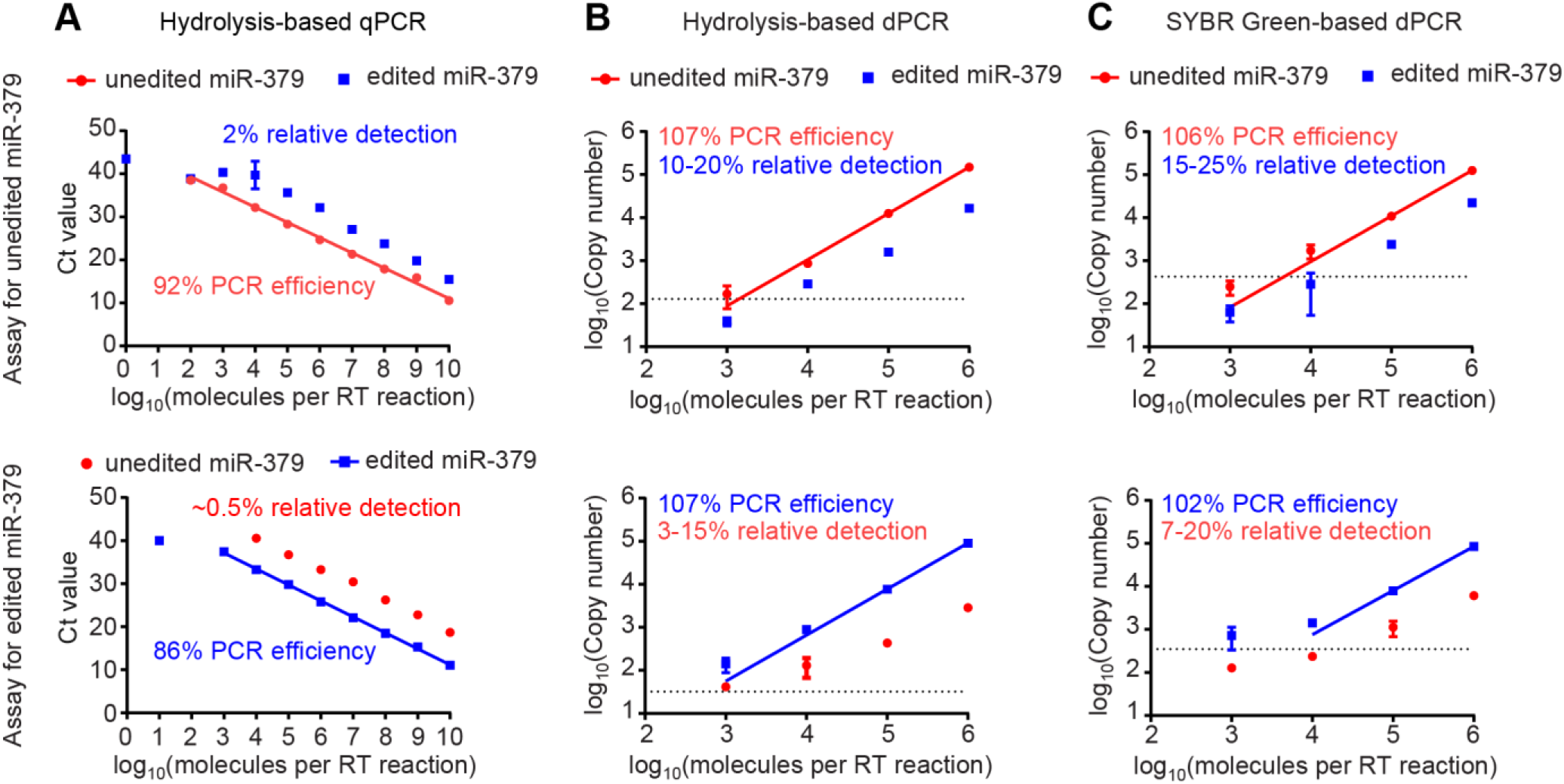
(**A**) Ct values for different dilutions of unedited (red circles) and edited (blue squares) miR-379 RNA oligonucleotides for unedited (top) and edited (bottom) miR-379 using hydrolysis probe-based qPCR. Error bars denote standard deviation; for points without error bars, the standard deviation was too small to be plotted. PCR efficiencies were calculated based on the slope. The average relative detection rate of non-target miR-379 was calculated based on the Ct difference to target miR-379. (**B**, **C**) Different dilutions of unedited (red circles) and edited (blue squares) miR-379 RNA for unedited (top) and edited (bottom) miR-379 quantified by hydrolysis probe-based dPCR (B) and (C) SYBR Green-based dPCR. Error bars denote standard deviation; for points without error bars, the standard deviation was too small to be plotted. The dotted line marks the background, i.e., the estimated “copy number” for negative control samples.

Unlike qPCR, dPCR separates the PCR reaction mixture into thousands of smaller compartments, either using droplets in an emulsion, or wells on a chip. This is done at a dilution that ensures that each compartment will either contain one single cDNA molecule or remain empty. After PCR, the number of positive partitions is determined (either using SYBR Green, or more commonly hydrolysis probes) and based on this, the number of cDNA molecules in the original mixture is calculated, yielding an absolute number of molecules without the need for a standard curve (Quan et al. 2018). Applying the two-tailed RT-qPCR method to chip-based dPCR showed a dynamic range up to 10^7^ molecules (Supplementary Figure 3), and satisfactory linearity with PCR efficiencies around 95–110% (Figure 3B and C, Supplementary Figure 3). Once again, relative detection of the non-target miR-379 was higher than previously, with 10– 20% detection of edited miR-379 with unedited miR-379-specific primers, and 3–15% detection of unedited miR-379 with edited miR-379-specific primers (Figure 3B). While SYBR Green was superior to hydrolysis probes in a qPCR setting, for dPCR, SYBR Green led to a further loss of specificity with 15–25% relative detection of edited miR-379 by unedited miR-379-specific dPCR and 7–20% relative detection of unedited miR-379 by edited miR-379- specific dPCR (Figure 3C). Importantly, both specificity and sensitivity of the editing-specific two-tailed assays even in the sub-optimal dPCR setting were still superior to currently available commercial assays (Figure 1).

### Quantification of miR-379 editing frequency in human tissues

We proceeded to test the performance of the editing-specific RT-qPCR method on a panel of tissue RNA samples, both to demonstrate usability of the method for a broad range of tissues, and to assess the miR-379 editing frequency in different human tissues. For this, we used SYBR Green-based qPCR, as it was the method with the highest specificity and sensitivity. All three primers (unedited miR-379, edited miR-379 and pan-miR-379) gave Ct values within the linear range of the standard curve for all tissues. Levels of miR-379 expression were highest in brain tissues and the adrenal gland (Figure 4A and Supplementary Figure 4A), which is in line with previous publications reporting high expression for the miR-379 cluster in the brain (Labialle et al. 2014). There was a strong correlation between levels of total miR-379 based on pan-miR-379 RT-qPCR and on the sum of unedited and edited miR-379 (Figure 4B), indicating that the pan-miR-379 assay reliably detects both isoforms.

**Figure 4.**
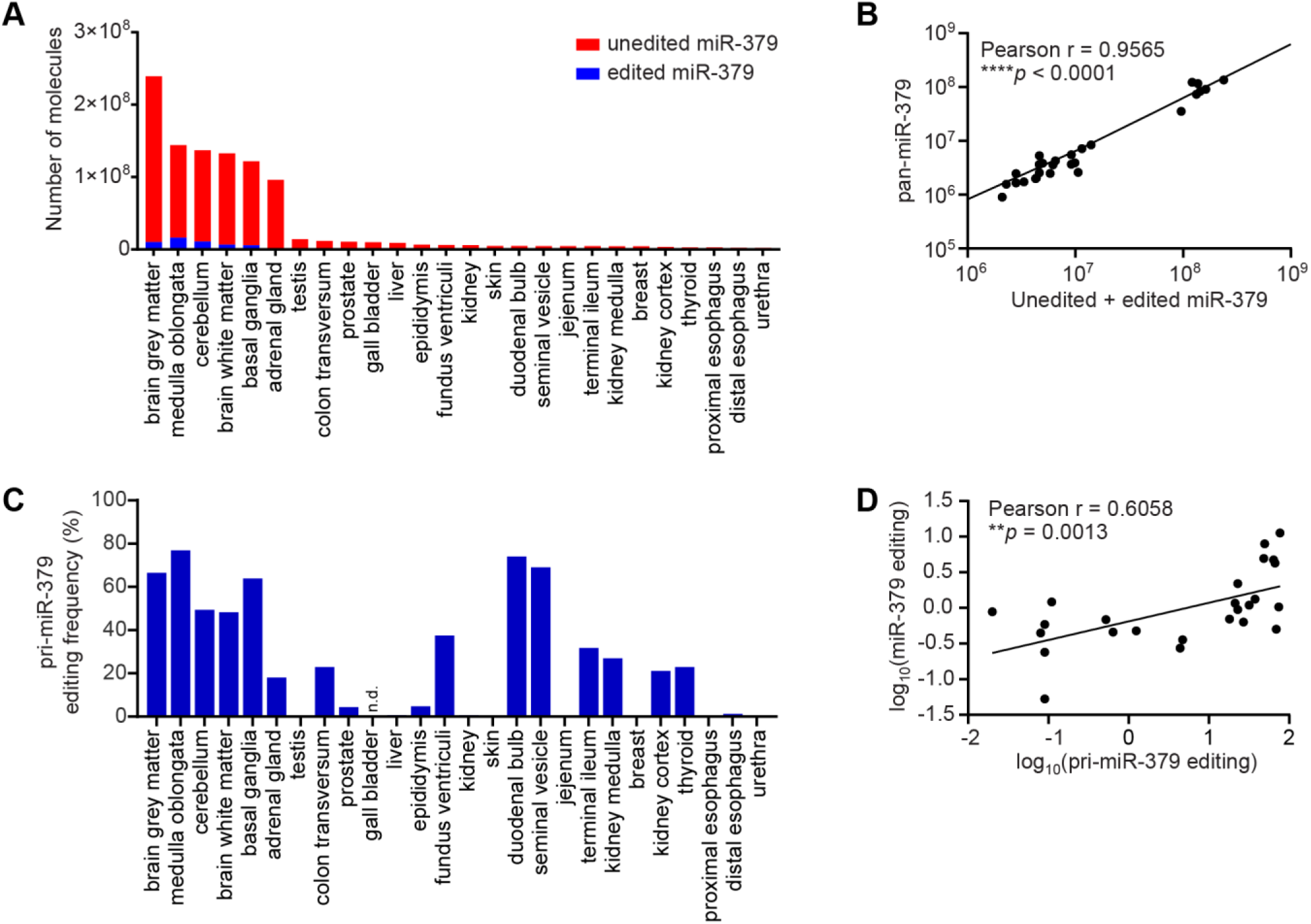
Expression and editing levels of miR-379 in a range of human tissues. (**A**) Number of molecules of unedited (red) and edited (blue) miR-379 in human tissues as determined by editing-specific two-tailed RT-qPCR. Absolute numbers were derived from standard curves of serially diluted RNA oligonucleotides, and normalized using the geometric mean of RNU24, RNU44, RNU48 and RNU66. (**B**) Correlation between the sum of unedited and edited miR-379 molecules and pan-miR-379 molecules. The *p* value was calculated by linear regression and Pearson correlation. (**C**) Editing frequencies of pri-miR-379 in a range of human tissues as determined by RT-PCR and Sanger sequencing. From gall bladder cDNA, pri-miR-379 could not be successfully PCR-amplified. n.d. = not detected. (**D**) Correlation between editing frequencies of mature miR-379 and pri-miR-379 in human tissues. Variables were log-transformed to enable meaningful linear regression and Pearson correlation.

Editing frequencies for tissues ranged between 0.24–11.2%, except for skin with a frequency of 0.05%. Based on the relative detection rates of the assays (Figure 2B), any editing frequencies between 0.08–99.6% should be correctly quantified by the assays. The detection of edited miR-379 is therefore likely not due to unspecific amplification of unedited miR-379 present in the sample, with the exception of the skin sample, in which edited miR-379 may not be present. The editing frequency of mature miR-379 was between 4–12% for brain tissues (previously estimated at 15% (Kawahara et al. 2008)), around 2% for the thyroid, and around 1% or lower for all other tissues (Supplementary Figure 4B).

Sanger sequencing of reverse-transcribed pri-miR-379 indicated higher editing frequencies across tissues (Figure 4C), which is expected, as pri-miR-379 editing is reported to reduce miR-379 maturation (Kawahara et al. 2008). Despite the discrepancy in magnitude, there was a statistically significant correlation between pri-miR-379 and mature miR-379 editing frequencies (Figure 4D). Editing frequencies of both pri-miR-379 and mature miR-379 were highest in the brain, in line with previous studies (Warnefors et al. 2014) and with reports that ADAR2 protein, which is responsible for miR-379 editing (Kawahara et al. 2008), is mainly expressed in the brain (Melcher et al. 1996, Seitz et al. 2004). Despite this, we did not find a strong correlation between miR-379 editing frequencies and levels of *ADAR* or *ADARB1* mRNAs, which code for ADAR1 and ADAR2 proteins respectively (Supplementary Figures 5 and 6). This supports reports that ADAR transcript levels do not always reflect editing activity (Wahlstedt et al. 2009).

### Quantification of miR-379 editing isoforms in a prostate cancer cohort

Finally, we assessed the clinical utility of our newly developed assays. We selected a clinical cohort of transurethral resections of the prostate to study miR-379 isoform regulation in PC. The cohort contained 23 tissue samples from patients with benign prostatic hyperplasia (BPH), and 47 samples from patients with PC.

We were able to detect both unedited and edited miR-379 in all samples (Supplementary Figure 7A), and the sum of the two isoforms correlated very well with the pan-miR-379 levels (Supplementary Figure 8). Editing frequencies were in the range of 0.77–52%, which is well within the previously defined reliable range of 0.08–99.6%. There were no significant differences in relative expression comparing BPH to PC samples with any of the three assays alone (Supplementary Figure 7A). However, the editing frequency of miR-379 was significantly higher in PC samples than BPH samples (Figure 5A). This indicates that quantifying both editing isoforms individually can reveal clinical information that cannot be detected by any single assay.

**Figure 5.**
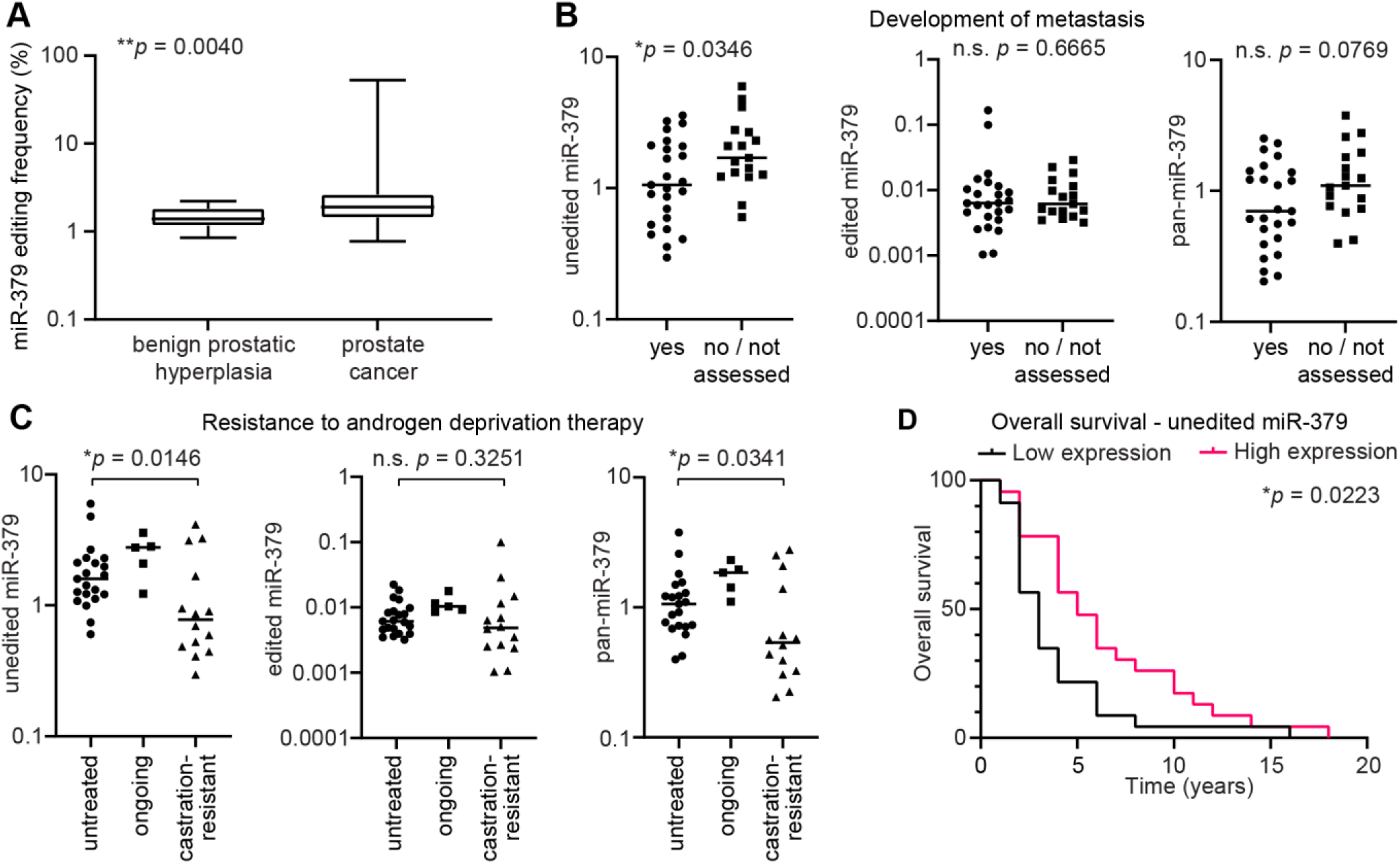
Deregulation of miR-379 editing and specific miR-379 isoforms in a prostate cancer cohort. (**A**) Editing frequency of miR-379 in 23 BPH and 47 PC patients calculated based on absolute numbers derived from standard curves. Box plot marks the median and upper and lower quartiles, whiskers denote the range of values. Exact *p* value was calculated using Mann-Whitney U test. (**B**) Comparison of relative expression of unedited miR-379 (left), edited miR-379 (middle) and pan-miR-379 (right) in PC patients that developed metastasis (n = 25) and those that did not get metastases or in which metastasis was not suspected and therefore not assessed (n = 17). Individual values and the median are shown. Exact *p* values were calculated using Mann-Whitney U tests. n.s. = not significant. (**C**) Comparison of relative expression of unedited miR-379 (left), edited miR-379 (middle) and pan-miR-379 (right) in hormone-naïve untreated PC patients (n = 21), those that were currently undergoing hormone treatment (n = 5), and those with castration-resistant PC (n = 14). Individual values and the median are shown. Exact *p* values were calculated using Mann-Whitney U tests. n.s. = not significant. (**D**) Survival analysis of patients with high or low relative expression of unedited miR-379. Patients were sorted by unedited miR-379 expression and divided into two groups at the median (n = 23 in each group). The *p* value was calculated using Log-rank test.

We then focused on the PC patients and compared the association of the expression of different miR-379 isoforms with certain clinical and pathological parameters. No clear expression differences of any miR-379 isoform were observed regarding tumour stage or histological grade (Supplementary Figure 7B and C). However, relative expression of unedited miR-379 was significantly lower in patients that already had or later developed metastases, whereas there were no statistically significant differences for edited miR-379 or pan-miR-379 (Figure 5B). This indicates that the predictive value of miR-379 only becomes apparent when performing isoform-specific quantification, and that edited miR-379 can be a confounding factor that masks deregulation when performing isoform-blind measurements. It is likely that other isoform-specific miRNA deregulation events with clinical potential have gone unnoticed due to the lack of suitable qPCR assays.

Furthermore, patients with castration-resistant disease following hormone deprivation treatment had significantly lower expression of unedited miR-379 and pan-miR-379, but not edited miR-379 (Figure 5C). We also performed survival analysis and found that patients with low levels of unedited miR-379 had significantly shorter overall survival than patients with high levels (Figure 5D). The same was observed for pan-miR-379, but not edited miR-379 (Supplementary Figure 7D). Unlike the comparison between BPH and PC, none of the comparisons within the PC patient group showed significant differences in miR-379 editing frequency (Supplementary Figure 9).

The fact that only unedited miR-379 was deregulated upon PC progression could suggest a potential role for this isoform in the suppression of castration resistance and metastasis, whereas edited miR-379 may not have the same role. A recent publication proposed opposite roles for unedited and edited miR-379 in multiple cancers (Xu et al. 2019). This study found unedited miR-379 to have a tumour-promoting role, whereas edited miR-379 inhibited cancer cell proliferation. However, the authors did not show any *in vitro* nor *in vivo* studies on PC cell lines, so that the role of miR-379 and its isoforms may be different in PC. This possibility is supported by a large-scale bioinformatic analysis of the TCGA dataset, which found that while miR-379 editing was reduced in most tumours compared to normal tissues, it was increased in prostate tumours compared to normal tissue (Pinto et al. 2017). This matches findings in other publications that expression of *ADARB1* mRNA is upregulated in PC, but downregulated in many other cancer types (Paz-Yaacov et al. 2015). These *in silico* studies support our findings of increased miR-379 editing in PC compared to BPH.

A limitation of our study lies in the rather small size of the analysed patient cohort, so it will be important to confirm our findings on similar cohorts in future studies. If it does hold true that unedited miR-379 is the only isoform associated with PC progression, and there is no evidence for a direct role of edited miR-379, the increase in miR-379 editing frequency in PC compared to BPH could serve as a mechanism to downregulate unedited miR-379 expression. Editing of pri-miR-379 inhibits the maturation of miR-379 (Kawahara et al. 2008), which is supported by the finding that miR-379 editing frequency was negatively correlated with total miR-379 expression in the analysed patient cohort (Supplementary Figure 10).

Of course, the discussed potential functional roles of the two miR-379 isoforms are mostly speculative at this point, and would need to be confirmed mechanistically in future studies. It is also possible that, rather than a driver event in tumour progression itself, the increased miR-379 editing frequency in PC is merely a consequence of increased ADAR2 activity. This does however not limit its potential use as a biomarker. Even if there is no functional role for miR-379 editing in PC, as long as ADAR2 activity is linked to relevant clinical parameters (Shaikhibrahim et al. 2013, Paz-Yaacov et al. 2015), a panel of editing-sensitive miRNA biomarkers such as miR-379 can be an easily accessible proxy for editing activity in the tumour. Using miRNA biomarkers as an indicator of editing activity can also be interesting for other diseases in which A-to-I editing has been shown to play a role, such as glioma (Maas et al. 2001), diabetes (Gan et al. 2006), amyotrophic lateral sclerosis (Hideyama et al. 2012), and chronic viral infections (Bass et al. 1989, Weiden et al. 2014).

In conclusion, we here describe the first validated qPCR technology that is able to distinguish A-to-I editing isoforms of an individual miRNA. The versatility of the described assays lies not only in their compatibility with different chemistries, but also in the ease with which the primers can be adapted to different sequences. The principle of selecting one hemiprobe to either cover the edited nucleotide (for editing-specific assays) or a non-editable part of the miRNA (for pan-miRNA analysis) is applicable to any miRNA, and thereby to virtually any disease in which miRNA biomarkers have clinical potential. Overall, we believe that the herein described development of A-to-I editing-specific RT-qPCR miRNA assays will serve as a useful tool for basic and translational research of miRNA function, and help develop better miRNA biomarkers for a range of diseases.

## Materials and Methods

### RNA and DNA oligonucleotides

RNA oligonucleotides of unedited miR-379, edited miR-379, miR-380, unedited miR-411, edited miR-411, and miR-758 were purchased from Integrated DNA Technologies, dissolved in IDTE buffer, pH 7.5 (Integrated DNA Technologies, Coralville, IA), and then diluted in 10-fold dilution series for RT-qPCR. For experiments with yeast RNA background, 100 ng total yeast RNA (#AM7118, Invitrogen, Thermo Scientific, Vilnius, Lithuania) was added during reverse transcription (RT) sample preparation. DNA oligonucleotides were purchased from Invitrogen. Two-tailed RT primers were designed based on a hairpin sequence published by Androvic *et al.* (Androvic et al. 2017) with hemiprobes designed to bind unedited or edited miR-379, and primer arms optimized to prevent the formation of unwanted secondary structures. Secondary structures of RT primers as well as secondary structures and dimers for qPCR primers were calculated using the OligoAnalyzer tool (Integrated DNA Technologies). ZEN/Iowa Black FQ double-quenched FAM-coupled miR-379 hydrolysis probe was designed using the PrimerQuest tool (Integrated DNA Technologies) and purchased from Integrated DNA Technologies. Primers for pri-miR-379 RT-PCR were based on those published by Kawahara *et al.* (Kawahara et al. 2008). All RNA and DNA oligonucleotide sequences are listed in Supplementary Table 1 and 2.

### Tissue and patient cohorts

The isolation of RNA from 26 human tissues was described previously (Lundwall et al. 2002). All tissues used in this study were obtained from patients undergoing surgery for neoplastic disease, or from autopsies.

Samples were collected from patients with voiding problems undergoing transurethral resection of the prostate (TURP) at Malmö University Hospital in 1990–1999. Small RNA isolation from formalin-fixed paraffin-embedded tissue sections using a modified procedure with the mirVana miRNA Isolation Kit (Ambion, Austin, TX) was previously described (Hagman et al. 2010). In this study, 23 patients with BPH and 47 patients with PC were included. The clinical characteristics of the cohort are summarised in Supplementary Table 3.

Ethical approval for the patient cohort was obtained from the Regional Ethical Review Board in Lund, and we adhered to the Helsinki declaration for all work with human tissues.

### Two-tailed RT-qPCR

For two-tailed RT-qPCR of miR-379, samples were reverse transcribed with the qScript Flex kit (#95049-100, Quantabio, Beverly, MA) using 2 μl 5x reaction mix, 1 μl GSP enhancer, 0.05 μM two-tailed RT primer, and 0.5 μl reverse transcriptase in a total volume of 10 μl. RT was performed at 25°C for 1 h, stopped at 85°C for 5 min and samples held at 4°C. RT products were used for qPCR or dPCR immediately. The input for RT was 1 μg RNA for the human tissue panel and 25 ng for patient samples.

The qPCR was performed with PowerUp SYBR Green Master Mix (#A25742, Thermo Scientific, Vilnius, Lithuania) using 400 nM forward and reverse primers (Supplementary Table 2). The RT product constituted up to 1/10^th^ of the total qPCR reaction volume. Samples were assayed in triplicates using the QuantStudio 7 Flex qPCR machine (Applied Biosystems). The qPCR program consisted of 30 s initial denaturation at 95°C, followed by 45 cycles of 30 s at 95°C and 20 s at 60°C. Melt curve analysis was performed to exclude the amplification of unspecific products. Absolute numbers of molecules for calculation of editing frequencies were interpolated from the Ct values by use of standard curves based on 10-fold dilutions of synthetic miR-379 oligonucleotides. For relative expression, the ΔCt method was used to normalize miR-379 Ct values to housekeeping small RNAs (see below).

For hydrolysis-based detection of miR-379 amplification, PrimeTime Gene Expression Master Mix (#1055772, Integrated DNA Technologies) was used with 400 nM primers and 250 nM hydrolysis probe. qPCR was performed in triplicates using a QuantStudio 7 Flex qPCR machine with initial denaturation at 95°C for 3 min, followed by 45 cycles of 5 s at 95°C and 30 s at 60°C.

### Two-tailed RT-dPCR

Digital PCR (dPCR) was carried out using QuantStudio 3D Digital PCR Master Mix v2 (Applied Biosystems, Frederick, MD), 400 nM primers, 1 μl cDNA per reaction, and either 250 nM hydrolysis probe or 2x SYBR Green I (Life Technologies, Eugene, OR). A volume of 14.5 μl reaction mix was loaded onto QuantStudio 3D Digital PCR 20K chips v2 (Applied Biosystems, Pleasanton, CA) in duplicates. Amplification was carried out with initial denaturation at 96°C for 10 min, and a total of 40 cycles of 2 min at 60°C alternated with 30 s intervals at 98°C. The chips were scanned on a QuantStudio 3D Digital PCR System (Applied Biosystems). Using QuantStudio 3D AnalysisSuite Cloud Software (Thermo Scientific), a threshold separating positive from negative wells was selected manually and applied to all chips to determine absolute copy numbers for each sample.

### RT-qPCR using commercial TaqMan assays

Reverse transcription with the TaqMan Advanced microRNA kit (#A28007, Applied Biosystems, Carlsbad, CA) was performed according to the manufacturer’s instructions. Briefly, the procedure consisted of adding a poly(A)tail and an adapter to the RNA, followed by reverse transcription using a universal RT primer and pre-amplification over 14 cycles. qPCR was then performed in quadruplicates using a TaqMan Advanced microRNA assay for miR-379 (#478077_mir) and TaqMan Fast Advanced Master Mix (#4444557, Applied Biosystems, Austin, TX) according to the manufacturer’s instructions on a QuantStudio 7 Flex machine.

For RT-qPCR of small RNA housekeeping controls, TaqMan microRNA assays (Applied Biosystems, Pleasanton, CA) were used according to the manufacturer’s instructions on 100 ng (human tissue panel) or 1 ng (patient cohort samples) of RNA in quadruplicates. For the human tissue panel, the geometric mean of RNU24 (#001001), RNU44 (#001094), RNU48 (#001006), and RNU66 (#001002) was used for normalization. For the patient cohort, the geometric mean of U47 (#001223), RNU48 and RNU66 was used for normalization. These combinations of housekeeping genes have previously been identified as optimal for these sample sets (Larne et al. 2013).

### Total cDNA synthesis

To prepare samples for pri-miR-379 and gene expression analysis, 2 μg total RNA was treated with DNase I (Thermo Scientific, Vilnius, Lithuania) according to the manufacturer’s instructions at 37°C for 30 min. Total cDNA was synthesised using the RevertAid H Minus First Strand cDNA Synthesis kit (#K1632, Thermo Scientific) according to the manufacturer’s instructions.

### Gene expression analysis

For gene expression analysis, cDNA was diluted 1:5 and 1 μl of the diluted cDNA was used for qPCR using TaqMan Gene expression assays and TaqMan Gene expression Master Mix (#4369016, Thermo Scientific) according to the manufacturer’s instructions. qPCR was performed in triplicates on a QuantStudio 7 Flex machine. Expression of *ADAR* (Hs00241666_m1) and *ADARB1* (Hs00953723_m1) was normalized to the geometric mean of *GUSB* (Hs99999908_m1), *PGK1* (Hs99999906_m1) and *GAPDH* (Hs02758991_g1) using the ΔCt method.

### PCR and Sanger sequencing of pri-miR-379

Pri-miR-379 cDNA was amplified from 5 μl of undiluted total cDNA with 1 U Phusion Hot Start II DNA polymerase (Thermo Scientific, Vilnius, Lithuania), 1x HF buffer, 200 μM dNTPs and 0.5 μM primers (Supplementary Table 2) in a total reaction volume of 50 μl. The reaction program consisted of 3 min initial denaturation at 98°C, followed by 35 cycles of denaturation at 98°C for 15 s, annealing at 64.7°C for 30 s and extension at 72°C for 30 s. After a final extension step for 10 min at 72°C, the PCR products were held at 4°C until PCR purification using the QIAquick PCR purification kit (#28106, Qiagen, Hilden, Germany). DNA concentrations were determined using NanoDrop 2000 (Thermo Scientific), and PCR products were sent for Sanger sequencing to Eurofins Genomics (Köln, Germany) using the forward primer used for the initial amplification. From the resulting chromatograms, peak heights of the individual nucleotides at the editing site were measured using SnapGene Viewer software (GSL Biotech; available at https://snapgene.com) to calculate relative editing frequencies.

### Statistical analysis

GraphPad Prism 9 (GraphPad software, La Jolla, CA) was used to interpolate absolute copy numbers from standard curves, and to perform all statistical analyses. PCR efficiencies were calculated from the slope of the standard curves as follows:

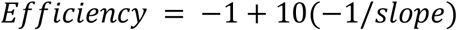

For association between variables, linear regression and Pearson correlation were used. If necessary, variables were log-transformed before regression and correlation analysis. For patient cohorts, non-parametric Mann-Whitney U tests were used to compare groups, and Log-rank test was used for survival analysis. A significance level of α = 0.05 was chosen for all statistical tests.

## Supplementary Data

SupplementaryFile1.pdf contains Supplementary Tables 1–4 and Supplementary Figures 1–10.

## Author contributions

GV and YC conceived and designed the study and wrote the manuscript. GV performed all experiments. AE and AB provided clinical samples and comments during the preparation of the manuscript.

## Acknowledgements

We are grateful to Prof. Dr. Åke Lundwall for making the human tissue panel available to us. For the prostate cancer cohort, we thank Elise Nilsson for tissue handling, and Dr. Olivia Larne for the RNA extraction. We are further grateful to Margareta Persson for excellent technical assistance.

