## Supplementary Material for "Quantification of microRNA editing using two-tailed RT-qPCR for improved biomarker discovery"

**Supplementary Table 1:** RNA oligonucleotide sequences.

| Name | Sequence |
|---|---|
| unedited miR-379-5p | 5’-UGGUAGACUAUGGAACGUAGG-3’ |
| edited miR-379-5p | 5’-UGGUIGACUAUGGAACGUAGG-3’ |
| miR-380-5p | 5’-UGGUUGACCAUAGAACAUGCGC-3’ |
| unedited miR-411-5p | 5’-UAGUAGACCGUAUAGCGUACG-3’ |
| edited miR-411-5p | 5’-UAGUIGACCGUAUAGCGUACG-3’ |
| miR-758-5p | 5’-GAUGGUUGACCAGAGAGCACAC-3’ |

**Supplementary Table 2:** DNA sequences of primers and hydrolysis probes

| Type | Name | Sequence |
|---|---|---|
| RT primer | unedited miR-379 RT | 5’-AGTCTATCATCATCAGAGCTAGAG AACCTAGCTCACCCACACTCCTAC-3’ |
| | edited miR-379 RT | 5’-AGTCCATCATCATCAGAGCTAGAG AACCTAGCTCACCCACACTCCTAC-3’ |
| | pan-miR-379 RT | 5’-ATAGTCTCATCATCAGAGCTAGAG AACCTAGCTCACCCACACTCCTAC-3’ |
| qPCR primer | unedited miR-379 fwd | 5’-GCAGTCTATCATCATCAGAGCTAG-3’ |
| | unedited miR-379 rev | 5’-GCGTGGTAGACTATGGAACG-3’ |
| | edited miR-379 fwd (for SYBR Green-based qPCR) | 5’-GCAGTCCATCATCATCAGAGCTAG-3’ |
| | edited miR-379 fwd_short (for hydrolysis-based qPCR) | 5’-AGTCCATCATCATCAGAGCTAG-3’ |
| | edited miR-379 rev | 5’-CGGGTGGACTATGGAACG-3’ |
| | pan-miR-379 fwd | 5’-GCATAGTCTCATCATCAGAGCTAG-3’ |
| | pan-miR-379 rev-1 (higher affinity for edited miR-379) | 5’-GCCTGGTGGACTATGGAACG-3’ |
| | pan-miR-379 rev-2 (higher affinity for unedited miR-379) | 5’-GCCTGGTAGACTATGGAACG-3’ |
| hydrolysis probe | miR-379 hydrolysis probe (binds both unedited and edited RT products) | 5’-(6-FAM)-AGGAGTGTG-(ZEN)-GGTGAGCTAGGTTCT-(Iowa Black FQ)-3’ |
| RT-PCR primer | pri-miR-379 fwd | 5’-TTTATGTTCACCATGTGCCTGC-3’ |
| | pri-miR-379 rev | 5’-TGCTGAAGCTAAACCACGTGTT-3’ |

**Supplementary Table 3:** Clinical characteristics of patients in the prostate cancer cohort. For PSA and age, the median and range are stated.

| | | Prostate cancer | Benign prostatic hyperplasia |
|---|---|---|---|
| **Total number of patients** | | 47 | 23 |
| **Age at TURP** | | 75 (63-89) years | 69 (56-89) years |
| **WHO Grade** | Grade I | 4 | - |
| | Grade II | 20 | - |
| | Grade III | 23 | - |
| **Clinical stage** | T1 | 9 | - |
| | T2 | 21 | - |
| | T3 | 13 | - |
| | T4 | 3 | - |
| | Data missing | 1 | |
| **PSA at surgery** | | 26.7 (0.2-672) ng/ml | 6.6 (3-19.4) ng/ml |
| **Metastasis** | Yes | 25 | - |
| | No | 8 | - |
| | Not suspected | 9 | - |
| | Data missing | 5 | - |

**Supplementary Table 4:** Different RT primer designs tested during the optimization of the assays. Different combinations for placement and length of the 5′ hemiprobe (red) and 3′ hemiprobe (blue) were tested in sequential experiments. The adenosine nucleotide subject to deamination upon RNA editing is highlighted in bold print. The RT primer that was selected for all subsequent experiments is highlighted with grey shading.

| Probe placement on miR-379 | Efficient amplification? | Specificity | Sensitivity |
|---|---|---|---|
| UGGUAGACUAUGGAACGUAGG | no | n/a | n/a |
| UGGUAGACUAUGGAACGUAGG | yes | < 1% | $10^3$ |
| UGGUAGACUAUGGAACGUAGG | yes | > 1% | n/a |
| UGGUAGACUAUGGAACGUAGG | no | n/a | n/a |
| UGGUAGACUAUGGAACGUAGG | yes | < 1% | $10^3$ |
| UGGUAGACUAUGGAACGUAGG | yes | < 1% | $10^3$ |
| UGGUAGACUAUGGAACGUAGG | no | n/a | n/a |
| UGGUAGACUAUGGAACGUAGG | yes | < 1% | $10^2$ |
| UGGUAGACUAUGGAACGUAGG | yes | > 1% | n/a |
| UGGUAGACUAUGGAACGUAGG | no | n/a | n/a |
| UGGUAGACUAUGGAACGUAGG | yes | < 1% | $10^2$ |
| UGGUAGACUAUGGAACGUAGG | yes | > 1% | n/a |

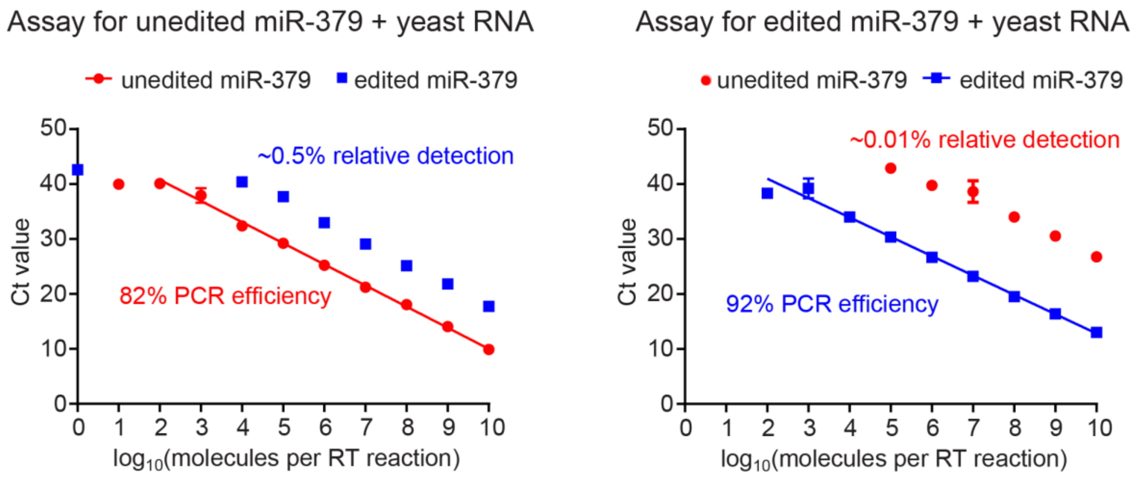


**Supplementary Figure 1.** Ct values for different dilutions of unedited (red circles) and edited (blue squares) miR-379 RNA oligonucleotides quantified by the two-tailed RT-qPCR assays specific for unedited (left) and edited (right) miR-379 with 100 ng yeast RNA contained in each reaction to create an artificial background. Error bars denote standard deviation; for points without error bars, the standard deviation was too small to be plotted. PCR efficiencies were calculated based on the slope. The average relative detection rate of non-target miR-379 was calculated based on the Ct difference to target miR-379.

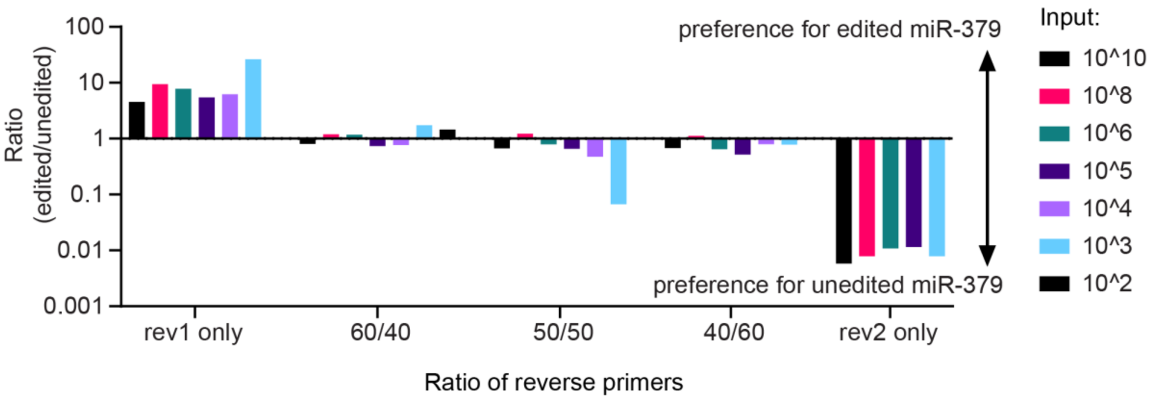


**Supplementary Figure 2.** Ratio between detection of different amounts of unedited and edited miR-379 by pan-miR-379 assays using different mixtures of reverse primers. A reverse primer ratio of 60/40 was deemed optimal as the detection ratios were closest to 1 with the least bias towards one or the other isoform, and this is the ratio that was subsequently used for all experiments in the study.

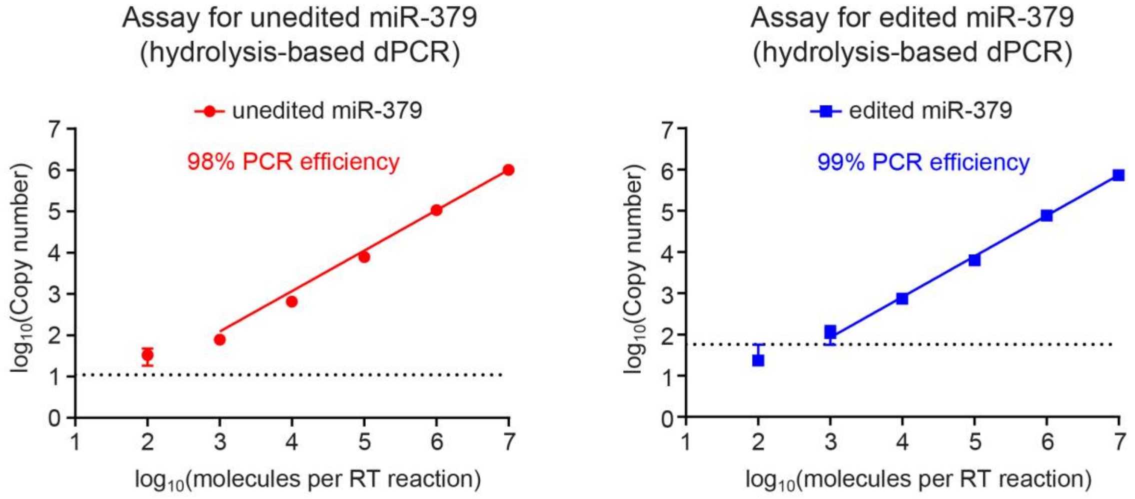


**Supplementary Figure 3.** Hydrolysis probe-based dPCR of different dilutions of unedited (left) and edited (right) miR-379 RNA oligonucleotides quantified by their specific two-tailed RT-qPCR assays, illustrating the dynamic range up to $10^7$ molecules input. Error bars denote standard deviation; for points without error bars, the standard deviation was too small to be plotted. PCR efficiencies were calculated based on the slope.

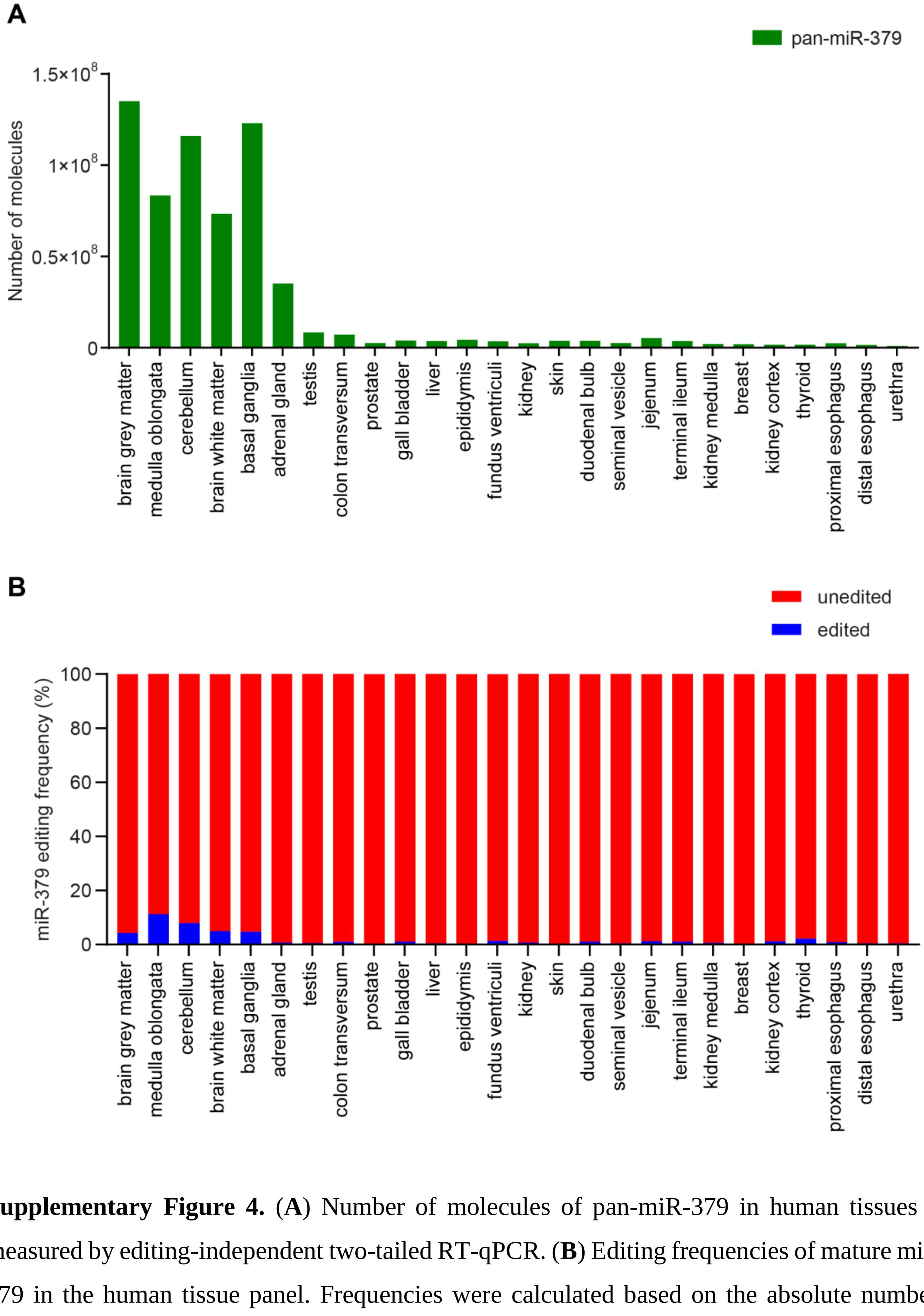


**Supplementary Figure 4.** (**A**) Number of molecules of pan-miR-379 in human tissues as measured by editing-independent two-tailed RT-qPCR. (**B**) Editing frequencies of mature miR-379 in the human tissue panel. Frequencies were calculated based on the absolute numbers presented in Figure 4A.

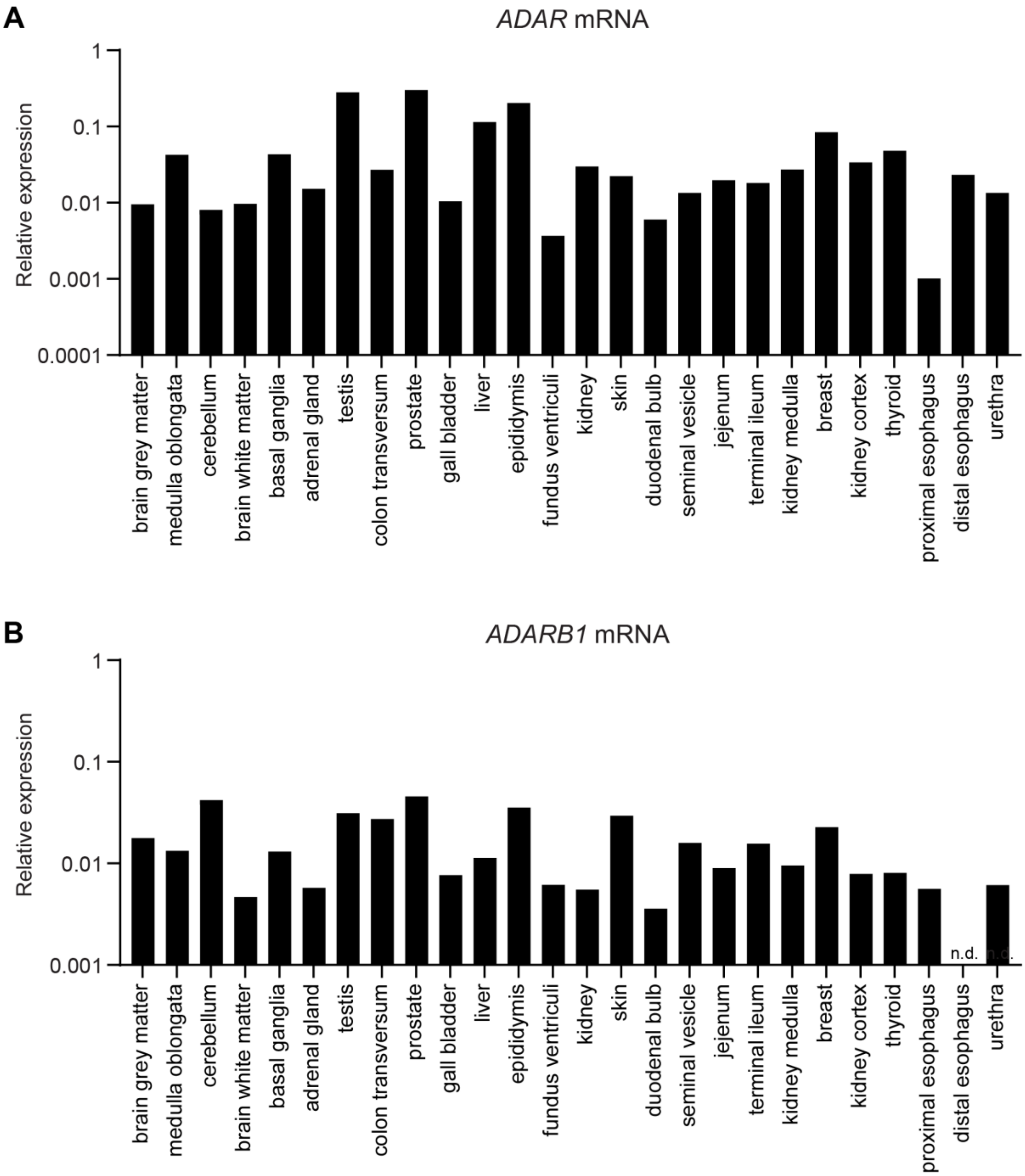


**Supplementary Figure 5.** (**A**) *ADAR* and (**B**) *ADARB1* mRNA expression in human tissues. In distal esophagus cDNA, *ADARB1* was undetectable. n.d. = not detected.

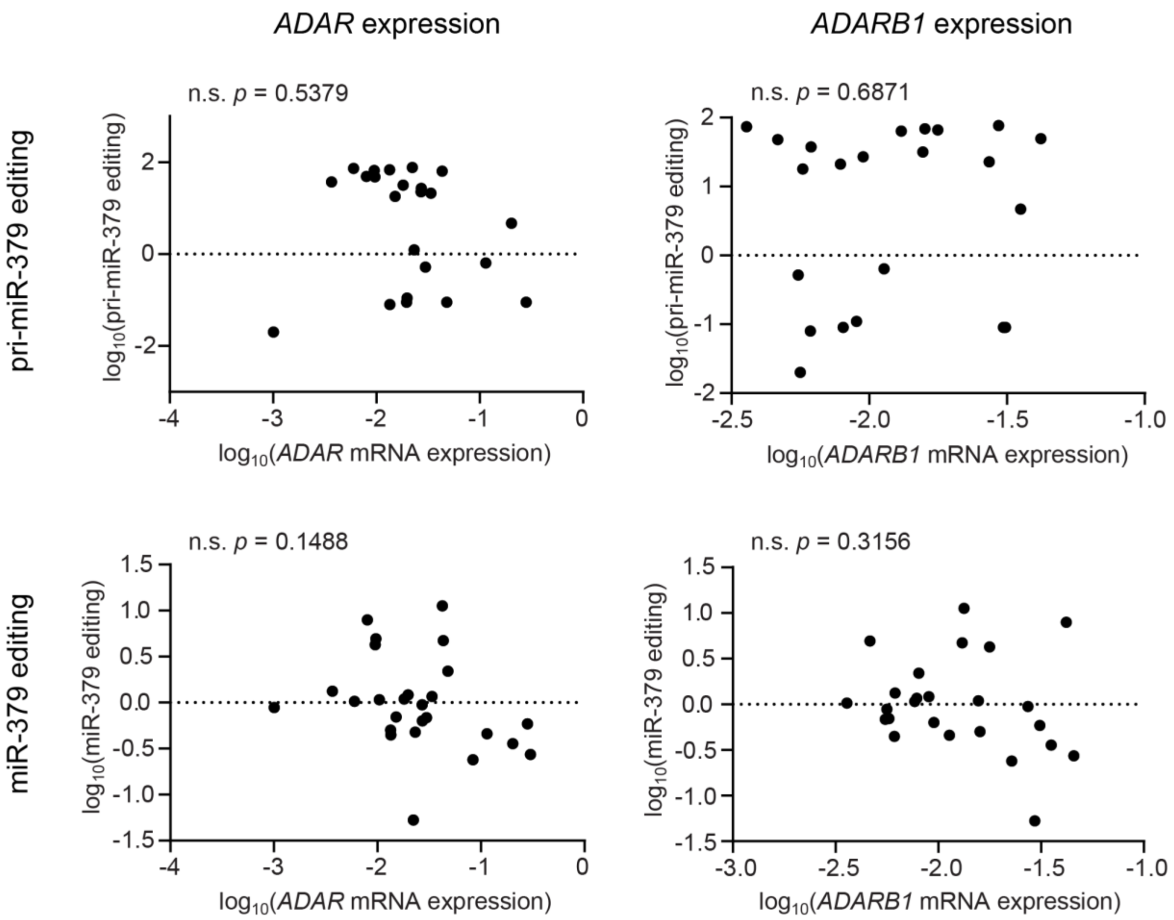


**Supplementary Figure 6.** Correlation analysis between pri-miR-379 (top) or miR-379 (bottom) editing frequencies and *ADAR* (left) or *ADARB1* (right) mRNA expression levels in human tissues. Variables were log-transformed to perform meaningful linear regression and Pearson correlation. n.s. = not significant.

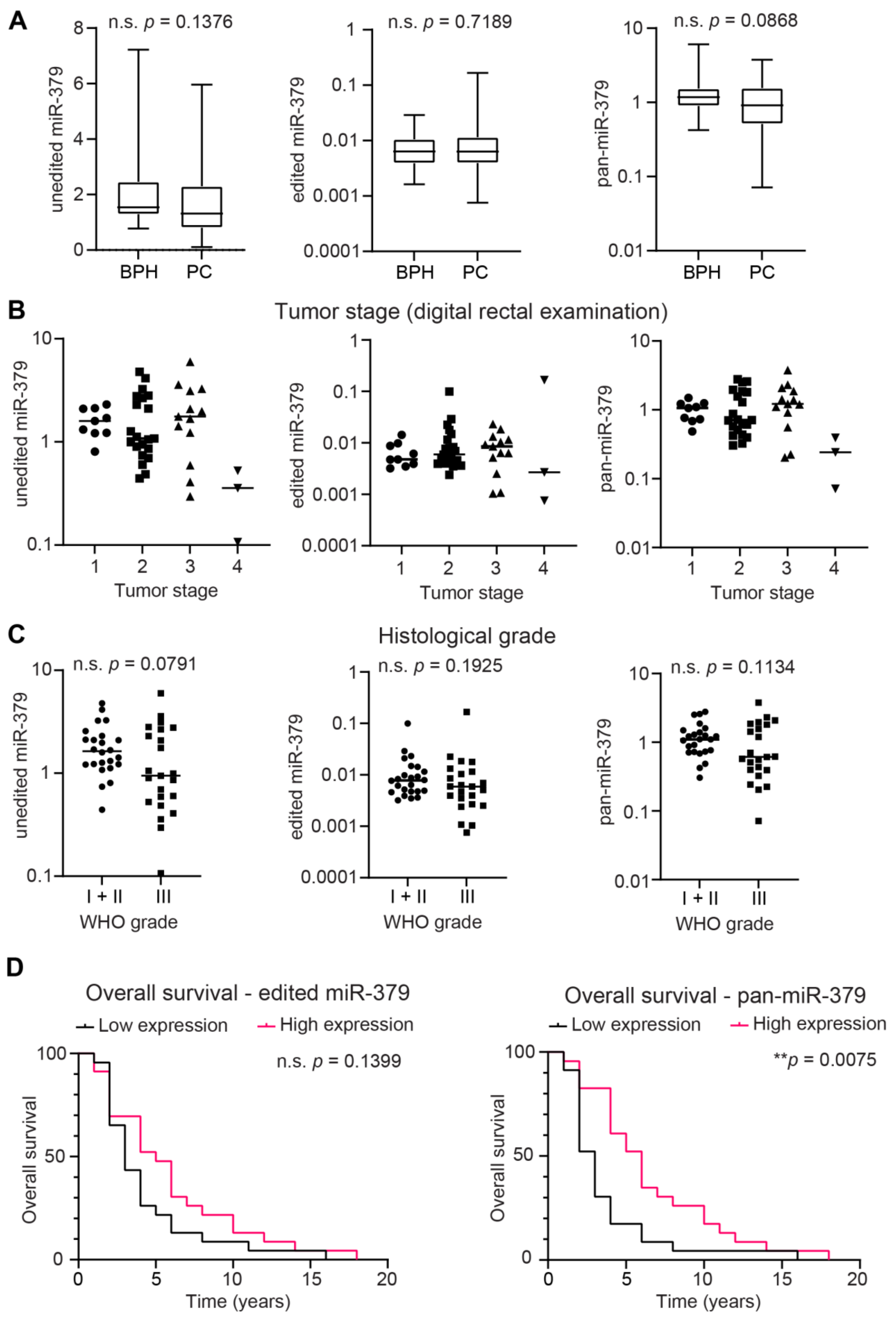


**Supplementary Figure 7.** (Legend on the following page)

**Supplementary Figure 7.** Expression of miR-379 isoforms in the PC patient cohort. (**A**) Relative expression of unedited miR-379 (left), edited miR-379 (middle) and pan-miR-379 (right) in patients with BPH (n = 23) and PC (n = 47). Box plot marks the median and upper and lower quartiles, whiskers denote the range of values. (**B**) Relative expression of unedited miR-379 (left), edited miR-379 (middle) and pan-miR-379 (right) by tumor stage (n = 9 for T1, n = 21 for T2, n = 13 for T3, n = 3 for T4). Individual values and the median are shown. (**C**) Relative expression of unedited miR-379 (left), edited miR-379 (middle) and pan-miR-379 (right) by pathological grade (n = 4 for Grade I, n = 20 for Grade II, n = 23 for Grade III). Individual values and the median are shown. (**D**) Survival analysis by edited miR-379 (left) and pan-miR-379 (right). Group comparisons were done by Mann-Whitney U tests. For survival analysis, patients were divided at the median by expression (n = 23 in each group), and curves were compared using log-rank test. n.s. = not significant.

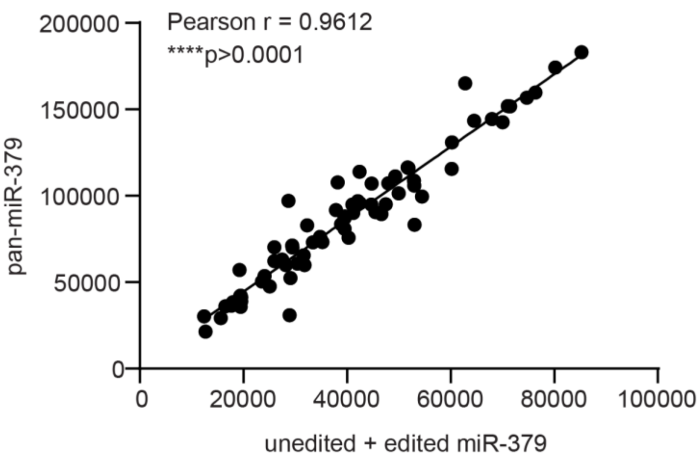


**Supplementary Figure 8** Correlation between the sum of unedited and edited miR-379 molecules and pan-miR-379 molecules (n = 70). The *p* value was calculated by linear regression and Pearson correlation.

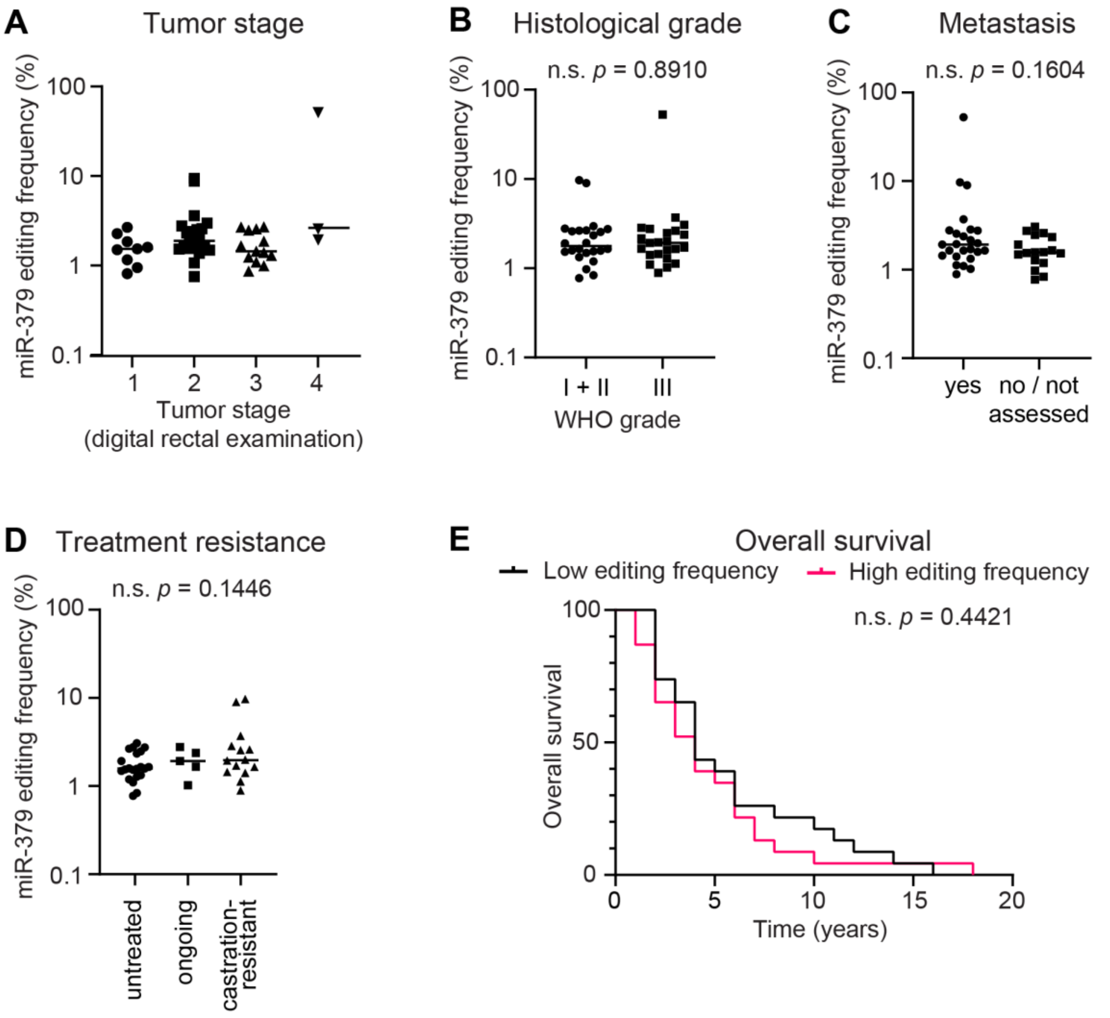


**Supplementary Figure 9.** Analyses of miR-379 editing frequency by (**A**) tumor stage (n = 9 for T1, n = 21 for T2, n = 13 for T3, n = 3 for T4), (**B**) histological grade (n = 4 for Grade I, n = 20 for Grade II, n = 23 for Grade III), (**C**) metastasis (n = 25 with metastasis, n = 17 without metastasis or not assessed), (**D**) androgen deprivation therapy (n = 14 hormone-naïve, n = 5 with ongoing hormone deprivation therapy, n = 21 castration-resistant), and (**E**) overall survival (n = 23 in each group). Individual values and the median are shown. Group comparisons were done by Mann-Whitney U tests. For survival analysis, patients were divided at the median by editing frequency, and curves were compared using log-rank test. n.s. = not significant.

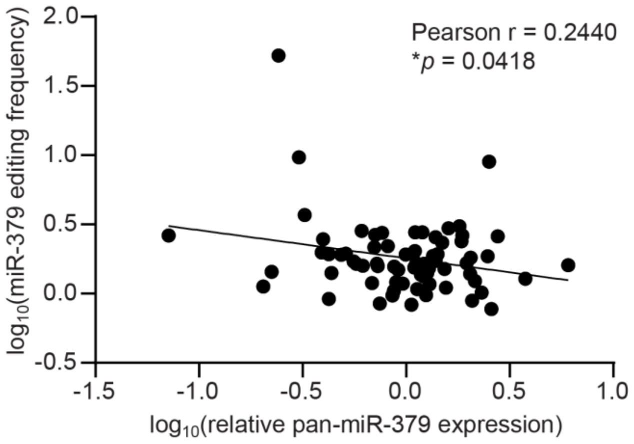


**Supplementary Figure 10.** Negative correlation between miR-379 editing frequency and relative pan-miR-379 expression in patient samples (n = 23). Both variables were log-transformed to enable meaningful linear regression and Pearson correlation.
